# A Congenic C57BL/6 *rd1* Mouse Model for Retinal Degeneration Research

**DOI:** 10.1101/2025.07.09.663548

**Authors:** Laurel C. Chandler, Apolonia Gardner, Constance L. Cepko

**Author notes:** Correspondence should be addressed to C.L.C., Department of Genetics, Harvard Medical School, 77 Avenue Louis Pasteur, Boston MA 02115, USA, Ph. These authors contributed equally: Laurel C. Chandler and Apolonia Gardner.

## Abstract

Retinitis pigmentosa is an inherited retinal disease caused by thousands of mutations in approximately 100 different genes. The most widely used mouse model for retinitis pigmentosa has the retinal degeneration 1 (*rd1*) mutation in the *Pde6b* gene, which elicits rapid retinal degeneration and vision loss. A major limitation of these models is that these *rd1* strains are not congenic, which prevents the use of appropriate controls. Furthermore, many strains have mutations in other genes which introduces genetic variability and may confound results. To address this issue, we backcrossed the *rd1* allele from FVB mice onto a C57BL/6J genetic background over many generations, producing a C57BL/6J.*Pde6b*^*rd1*^ strain that was confirmed to be congenic to C57BL/6J mice. We show that this strain recapitulates the electroretinogram and optomotor results expected for mouse strains containing the *rd1* mutation. Examination of retinal structure in cross sections of eyes isolated from C57BL/6J.*Pde6b*^*rd1*^ mice show a degree of thinning of the outer nuclear layer expected for a *rd1* mutation, resulting in nearly complete loss of the outer nuclear layer by postnatal day 35. We anticipate that this C57BL/6J.*Pde6b*^*rd1*^ strain could become an asset for the field of retinitis pigmentosa research.

---

Retinitis pigmentosa (RP) is the most common cause of inherited retinal degeneration, affecting ∼2 million people globally.^1^ The disease manifests as an initial loss of rods, affecting night vision, followed by the gradual degeneration of cones, which affects daylight vision.^2^ This disease can be caused by a wide variety of mutations in any of ∼100 genes linked to RP.^3^ The PDE6B gene, encoding for the phosphodiesterase 6 beta subunit, plays a crucial role in the light-responsive signal transduction pathway in rod photoreceptors.^4^ To model RP in mice, there are several strains with mutations in *Pde6b*, including those which carry the retinal degeneration 1 (*rd1*) allele, which leads to rapid retinal degeneration and vision loss. The *rd1* mutation includes an intronic murine leukemia viral insertion in exon 1 and a nonsense point mutation in exon 7, which together lead to complete loss of PDE6β protein.^5,6^ As the first and best-studied retinal degeneration mouse model, mice exhibiting the *rd1* mutation are widely used in RP research,^7^ with >1,000 articles published using this allele (PubMed search term “rd1”). However, strains which contain this mutation, including FVB, C3H, CBA, and SJL, have mutations in other genes, which may confound results. It is difficult to include the appropriate control strain as there is no congenic *rd1* mouse model, unlike other RP mouse strains such as *rd10* and P23H.^8^ In order to generate a congenic *rd1* mouse, we backcrossed FVB mice, which carry the *rd1* mutation, with C57BL/6J mice for many generations until the *rd1* allele was present on a C57BL/6J background. This study was approved by the Institutional Animal Care and Use Committee at Harvard University. C57BL/6J mice (strain #000664) were purchased from The Jackson Laboratory and FVB mice (strain #207) were purchased from Charles River Laboratories. The Jackson Laboratory’s SNP Genome Scanning service was used to investigate the genetic background of five mice from these backcrosses. These assays revealed that, of the SNPs assayed, all C57BL/6J.*Pde6b*^*rd1*^ animals had C57BL/6J alleles and 0% had the alleles of the FVB strain (Table 1).

**Table 1.** Determination of the background identity of the C57BL/6.*Pde6brd1* strain.

| Sample | FVB/NJ (% identity) | C57BL/6J (% identity) |
| --- | --- | --- |
| C57BL/6J.*Pde6b*<sup>*rd1*</sup> #1 | 0.00 | 100.00 |
| C57BL/6J.*Pde6b*<sup>*rd1*</sup> #2 | 0.00 | 100.00 |
| C57BL/6J.*Pde6b*<sup>*rd1*</sup> #3 | 0.00 | 100.00 |
| C57BL/6J.*Pde6b*<sup>*rd1*</sup> #4 | 0.00 | 100.00 |
| C57BL/6J.*Pde6b*<sup>*rd1*</sup> #5 | 0.00 | 100.00 |
| C57BL/6J control | 0.00 | 100.00 |
| FVB/NJ control | 100.00 | 0.00 |
| C57BL/6J x FVB/NJ heterozygous | 50.00 | 50.00 |

To characterize visual function in the C57BL/6J.*Pde6b*^*rd1*^ mice, electroretinogram (ERG) analysis was performed. To measure rod and cone function, scotopic and photopic ERGs were measured respectively using an Espion E3 System (Diagnosys LLC) as previously described.^9^ All animals were dark adapted for at least four hours prior to recording. Mice were anesthetized by intraperitoneal injection of ketamine and xylazine (100 and 10 mg/kg, respectively).

Tropicamide 1% eye solution (Bausch + Lomb) was used to dilate the pupil. Dim flashes were applied to elicit a scotopic ERG response at a 0.1 candela (cd) s/m^2^ intensity. The scotopic b-wave response was assessed in both wildtype C57BL/6J and C57BL/6J.*Pde6b*^*rd1*^ mice at postnatal day 20 (P20) in low light conditions to isolate the rod photoreceptors and their downstream connection to ON-bipolar cells. There was a statistically significant loss of scotopic ERG response in C57BL/6J.*Pde6b*^*rd1*^ compared to C57BL/6J mice (Figure 1A). All animals were then light adapted for six minutes with a white light background of 30 cd s/m^2^. Multiple white flashes were applied to elicit photopic ERG responses at 1 (peak), 10 (peak), 100 (xenon), and 1000 (xenon) cd s/m^2^ intensities with a background light of 30 cd s/m^2^. At all intensities, the photopic b-wave, which primarily measures the response from cone ON-bipolar cells, was significantly reduced in C57BL/6J.*Pde6b*^*rd1*^ compared to C57BL/6J mice (Figure 1B and C).

**Figure 1.**
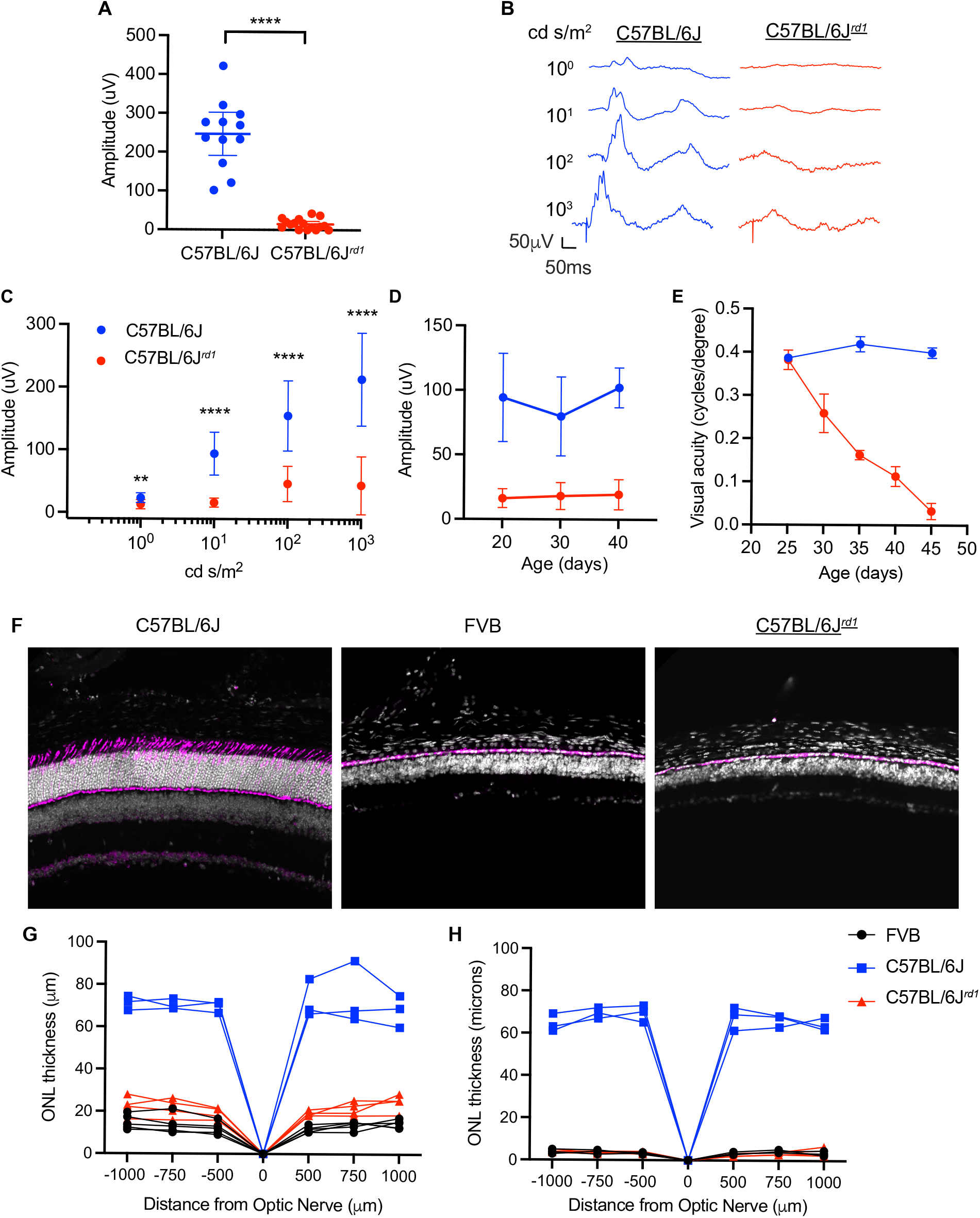
Examination of the visual function, visual acuity, and outer nuclear layer (ONL) thickness in C57BL/6J mice with the *rd1* mutation (C57BL/6J*rd1*) compared to those of the FVB strain. (A) Dark-adapted scotopic response at a 0.1 cd s/m2 flash intensity in P20 C57BL/6J (n=12) and C57BL/6J*rd1* (n=16) mice (unpaired T-test). (B) Representative light-adapted electroretinogram (ERG) traces in P20 C57BL/6J or C57BL/6J.*Pde6brd1* (C57BL/6Jrd1) mice subject to a range of flash intensities. (C) Light-adapted photopic b-wave in P20 mice at a range of flash intensities (n=12). ^**^p<0.01, ^****^p<0.0001 (two-way ANOVA with Šidak^’^s multiple comparison test). (D) Photopic b-wave at a flash intensity of 100 cd s/m2 over time at ages P20, 30 and 40 (n=12). (E) Visual acuity of wildtype C57BL/6J compared to C57BL/6J*rd1* mice as measured by the number of cycles/degree using the optomotor assay. (F) Representative retinal sections from wildtype C57BL/6J, FVB and C57BL/6J*rd1* mice at P20. Sections are stained with cone arrestin (magenta) and DAPI (white). ONL thickness in (G) P15 and (H) P35 mice measured using the DAPI channel at intervals along the dorsal-ventral axis with the most dorsal and ventral points at 1000 sssm from the optic nerve.

There was no further reduction in the b-wave amplitude when assessing the ERG response at 10 cd s/m^2^ at P30 and 40, suggesting that most of the response had already been lost by P20 (Figure 1D).

Although the ERG response is rapidly lost in RP mouse models, the optomotor response can be retained for significantly longer. The optomotor response was measured using the OptoMotry System (CerebralMechanics) under photopic conditions with a background luminance of ∼70 cd s/m^2^ as previously described.^10^ The contrast of the black and white stripes were displayed at 100% and the spatial frequency was set to 1.5 Hz. To assess visual acuity, the program changed the frequency of the stripes (cycles/degree) and the direction of the rotation (clockwise vs counterclockwise), while the examiner recorded whether head tracking was present. This was continued until the program had reached a threshold of acuity for each animal. Visual acuity was assessed every five days from P25-45 in C57BL/6J and C57BL/6J.*Pde6b*^*rd1*^ mice. At P25, there was no significant difference between the visual acuity of the C57BL/6J and C57BL/6J.*Pde6b*^*rd1*^ mice, however from P30 onwards there was a gradual loss of vision in the C57BL/6J.*Pde6b*^*rd1*^ mice while wildtype mice remained unchanged (Figure 1E).

To visualize retinal morphology in the C57BL/6J.*Pde6b*^*rd1*^ mice compared to FVB and C57BL/6J controls, eyes were harvested from mice and prepared for sectioning as previously described.^2^ Sections were cut at a thickness of 25 μm on a CM 3050S cryostat (Leica Microsystems) prior to staining with cone arrestin (1:3000 dilution, Millipore Sigma #AB15282) in blocking buffer (4% donkey serum, 0.3% Triton X-100, 0.3% bovine serum albumin) overnight at 4°C in a humidified chamber. The following day, sections were washed three times in PBS and incubated with anti-rabbit 647 secondary antibody (1:750, Jackson ImmunoResearch) in blocking buffer for two hours at room temperature. After staining, samples were rinsed three times in PBS and stained with DAPI for five minutes prior to three final washes in PBS. Samples were mounted with Fluoromount G (Southern Biotech) under coverslips and imaged on a Nikon Ti inverted microscope with a W1 Yokogawa Spinning disk with 50 μm pinhole disk and a Plan Apo λ 20×/0.75 DIC I objective. Representative eyecup sections at P20 for each of the three mouse strains showed the outer nuclear layer (ONL), which is visualized by cone arrestin staining, to be comparable in the C57BL/6J.*Pde6b*^*rd1*^ and FVB mouse strains, both of which were significantly thinner than the wildtype C57BL/6J controls (Figure 1F) suggesting substantial retinal degeneration by this age.

ONL thickness was also measured at set distances from the optic nerve along the dorsal-ventral axis. After sectioning, samples were stained with DAPI in staining buffer (1% Triton X-100, 4% normal donkey serum in PBS) for 2 hours at room temperature. After staining, slides were rinsed in PBS, mounted onto a coverslip and imaged on an Olympus VS200 Slide Scanner with a UPlan X Apo 10x/0.4 Air objective. Images were analyzed using OlyVIA software. The thickness of the ONL was measured at 500, 750, and 1000 μm intervals both dorsal and ventral of the optic nerve. There was a significant reduction in ONL thickness in both FVB and C57BL/6J.*Pde6b*^*rd1*^ mice at P15 relative to C57BL/6J controls (Figure 1G). This thickness was further reduced to ∼1 layer of nuclei in the FVB and C57BL/6J.*Pde6b*^*rd1*^ mice at age P35, whereas C57BL/6J controls retained normal ONL thicknesses (Figure 1H).

In this study, we generated a congenic mouse line carrying the *Pde6b*^*rd1*^ allele. This allele was transferred onto the C57BL/6J background in-house by backcrossing the FVB *rd1* mutation onto the C57BL/6J mouse background. After confirming that this new C57BL/6J.*Pde6b*^*rd1*^ strain has the background of the C57BL/6J strain, with no detectable alleles from the FVB strain, we performed both functional and structural assays to determine the extent and kinetics of degeneration in this mouse line. ERG analyses showed a significant reduction in scotopic (rod-mediated) and photopic (cone-mediated) vision by P20 that didn’t undergo further reduction at later ages, indicating that the C57BL/6J.*Pde6b*^*rd1*^ strain had lost most visual function by weaning age. We also performed an optomotor assay to assess visual acuity from P25-P45, which showed a steady decline in the C57BL/6J.*Pde6b*^*rd1*^ strain until a near total loss was achieved by P45. Consistent with the loss of visual function, ONL thickness was markedly reduced in the C57BL/6J.*Pde6b*^*rd1*^ strain relative to wildtype C57BL/6J controls at P15. By P35, the ONL was reduced to ∼1 nuclei layer in the C57BL/6.*Pde6b*^*rd1*^ strain. ONL thickness was reduced to approximately the same extent in FVB and C57BL/6.*Pde6b*^*rd1*^ at both timepoints suggesting that retention of the *rd1* allele from the FVB strain is sufficient for retinal degeneration.

Although there are other congenic/coisogenic strains of *Pde6b* mutant mice now available, including those generated by Chirinskaite and colleagues using CRISPR-Cas9 KO technology^11^ and another by The Jackson Laboratory induced via a chemically induced mutation (strain #004766), sequencing demonstrates that these other mutant strains contain disruptions additional to the *rd1* mutation and have slower kinetics and extents of degeneration.^11^ The C57BL/6J.*Pde6b*^*rd1*^ strain described in this study optimally combines the benefits of a congenic mouse model with rapid-onset retinal degeneration.

## Acknowledgments

The authors wish to thank Lucas Lin and Grace Wallick for their technical assistance. The authors wish to thank the Micron Core (Dr. Paula Montero-Llopis, Dr. Praju Vikas Anekal, and Dr. Adrienne Wells) for their assistance with microscopy. This study was funded by the Howard Hughes Medical Institute, the Harvard Brain Institute (L.C.C.), and the National Eye Institute (NIH F31 Ruth L. Kirschstein NRSA awarded to A.G.).

